# *Bacillus thuringiensis* bioinsecticide influences *Drosophila* oviposition decision

**DOI:** 10.1101/2023.03.07.531532

**Authors:** Aurélie Babin, Jean-Luc Gatti, Marylène Poirié

## Abstract

Behavioural avoidance has obvious benefits for animals facing environmental stressors such as pathogen-contaminated foods. Most current bioinsecticides are based on the environmental and opportunistic bacterium *Bacillus thuringiensis* (*Bt*) that kills targeted insect pests upon ingestion. While food and oviposition avoidance of *Bt* bioinsecticide by targeted insect species was reported, this remained to be addressed in non-target organisms, especially those affected by chronic exposure to *Bt* bioinsecticide such as *Drosophila* species. Here, using a two-choice oviposition test, we showed that female flies of three *Drosophila* species (four strains of *D. melanogaster*, *D. busckii* and *D. suzukii*) avoided laying eggs in the presence of *Bt* var. *kurstaki* bioinsecticide, with potential benefits for the offspring and female’s fitness. Avoidance occurred rapidly, regardless of the fraction of the bioinsecticide suspension (spores and toxin crystals versus soluble toxins/components) and independently of the female motivation for egg laying. Our results suggest that, in addition to recent findings of developmental and physiological alterations upon chronic exposure of non-target *Drosophila*, this bioinsecticide may have greater ecological implications in the field for the *Drosophila* community and their associated natural enemies than previously thought.

## 1. Introduction

When exposed to environmental stressors, animals face two main options: dealing with the stressor, which may ultimately lead to the evolution of special features, or physically avoiding it. In interactions with opportunistic pathogens, broad-sense immunity includes components for dealing with pathogens when interactions occur (physical barriers and cellular and humoral effectors of the immune system) as well as a behavioural component to physically avoid pathogens and reduce the infection risk.^[1–3]^ The immune response being costly (energy, nutrients, and immunopathology resulting from damage to host tissues by effectors of its innate immune response)^[2,4]^, obvious benefits come from physically avoid pathogens.

Behavioural avoidance of toxic compounds and microorganisms in a foraging context is well documented. Both innate avoidance (‘disgust’) and learned avoidance based on associative learning of hazardous food, are commonly expressed by vertebrates^[5]^ and invertebrates, mainly insects.^[6–8]^ For instance, phytophagous insects avoid plants that accumulate toxic alkaloids^[9]^ and the nematode *Caenorhabditis elegans* prefers feeding on non-pathogenic bacteria rather than pathogenic ones^[10,11]^. Exposed to opportunistic pathogens through their diet of overripe fruits, *Drosophila melanogaster* females are able to learn to adjust their preference for a food odour when that odour has previously been associated with a gut infection by the virulent bacterium *Pseudomonas entomophila*,^[12]^ as do *C. elegans* nematodes when exposed to pathogenic bacteria.^[13]^ *Drosophila melanogaster* males and females also express strong innate aversive responses to bacterial lipopolysaccharides when feeding and egg laying respectively, mediated by dTRPA1 cation channels of gustatory neurons.^[14]^

Naturally ubiquitous in the environment, *Bacillus thuringiensis* (*Bt*) is an opportunistic Gram-positive bacterium, which synthesizes insecticidal toxins including Cry proteins as crystals along with spores.^[15,16]^ The insecticidal action relies on the organisms’ feeding activity on *Bt*-contaminated food sources.^[17]^ In the context of the growing global food demand and the need for safer and more specific insect pest control, these natural insecticidal properties have led the development of *Bt-*based bioinsecticides (products made of viable *Bt* spores and toxin crystals) or *Bt* transgenic crops to control insect pests in agriculture and forestry (mainly Lepidoptera), and mosquitoes and black flies (Diptera).^[18,19]^ Many studies concluded that *Bt* bioinsecticides and *Bt* crops are harmless or have limited impacts on the non-target fauna.^[20,21]^ However, the partial targeting specificity of Cry toxins and the potential for environmental accumulation of spores and toxins upon repeated treatments have raised concern about potential side-effects on non-target organisms.^[16,22–25]^ In insects, recent studies have reported deleterious effects of the Lepidoptera-targeting *Bt* var. *kurstaki* (*Btk*) bioinsecticide on several species of non-target *Drosophila* flies likely present in *Btk*-treated areas. Chronic exposure of fly larvae to subacute doses through the diet altered their growth, development duration, survival, and complete development success.^[26–29]^ *Btk* bioinsecticide also impacted the larval metabolism and midgut physiology, impairing protein digestion and disturbing the gut epithelium organisation.^[28]^ One way for non-target insects that would alleviate *Bt* bioinsecticide impacts is the expression of behavioural avoidance of *Bt*-treated substrates. As *Bt* bioinsecticides act after ingestion, behavioural avoidance would be advantageous upon food foraging, but also upon female oviposition with direct benefits for the offspring and indirect benefits for the female’s fitness.

So far, *Bt* behavioural avoidance has been investigated mainly in *Bt*-target invertebrates: studies have reported no change in the oviposition behaviour of *Culex* mosquitoes exposed to *Bt* var. *israelensis*^[30]^ or in the feeding behaviour of the Western corn rootworm *Diabrotica virgifera virgifera*,^[31]^ and even an attractive effect of *Bt* maize on the oviposition of the fall armyworm *Spodoptera frugiperda*.^[32]^ By contrast, behavioural avoidance of *Bt* upon food foraging was reported in the nematode *Caenorhabditis elegans*^[33–36]^ and in two Lepidopteran pests, the cotton bollworm *Helicoverpa armigera* and the cotton leafworm *Spodoptera litura*.^[37]^ Females of *H. armigera* and of the diamondback moth *Plutella xylostella* also avoid *Bt* when laying eggs in a choice situation.^[38,39]^ *Bt* avoidance was also reported in insects’ offspring: neonates of the European corn borer*, Ostrinia nubilalis*, disperse more on *Bt* corn^[40]^ and avoid *Bt* when facing a choice with untreated diet,^[41]^ while neonates of the tobacco budworm *Heliothis virescens* avoid diets containing Cry toxins or the *Bt* bioinsecticide at doses that do not alter their development and survival.^[42]^

By contrast, *Bt* behavioural avoidance has been scarcely addressed in non-target invertebrates. Foraging activity and learning ability of *Apis mellifera ligustica* honey bees remained unchanged on *Bt* corn,^[43]^ while collective nest building and prey attacks were altered by cuticular *Bt* inoculation to the African social spider *Stegodyphus dumicola*.^[44]^ Altered reproduction and survival were recorded in *Bombus terrestris* bumble bees exposed to *Bt* depending on the *Bt* subspecies and the exposure route, but without altering the foraging behaviour and colony performance.^[45]^ *Bt* bioinsecticides being increasingly applied in the field, studies exploring the behavioural avoidance by non-target invertebrates are needed for an accurate assessment of the potential bioinsecticide side-effects on non-target fauna.

Here, we explored the expression of behavioural avoidance toward the lepidopteran-targeting *Bt* var. *kurstaki* (*Btk*) bioinsecticide by non-target *Drosophila* species that exhibit developmental and physiological alterations in the chronic presence of bioinsecticide.^[27,28]^ *Drosophila* larvae are particularly exposed to food-borne stressors as they intensively search for food to fuel their exponential growth but have a low dispersal capacity. Bioinsecticide avoidance by adult females when searching for oviposition sites would mitigate the consequences on larval development. We focused on three *Drosophila* species with different ecological features and varying developmental alterations elicited by chronic *Btk* exposure: two cosmopolitan domestic species which frequently coexist on overripe fruits, *D*. *melanogaster* (four strains) and the phylogenetically distant and opportunistic *D. busckii*,^[46–50]^ and the invasive *D. suzukii* that feeds and lays eggs on ripe fruits and is a threat to agriculture.^[51–54]^ We measured the females’ oviposition preference in two-choice tests where they were offered food with or without *Btk* bioinsecticide at a specific dose. The preference dynamics during the choice test was recorded and the effect of different fractions of the *Btk* bioinsecticide suspension (spores and toxin crystals, and soluble toxins/components) on the fly preference was also assessed.

## 2. Material and Methods

### (a) Fly stocks

Four *D. melanogaster* strains were tested: the wild-type Canton-S (Bloomington Drosophila Center) used here as a reference strain, the wild-type “Nasrallah” from Tunisia (strain 1333, Gif-sur-Yvette), a wild-type strain “Sefra” derived from flies collected in Southern France in 2013, and the yellow-white double mutant YW1118 (gift from Dr. B. Charroux, IBD, Marseille-Luminy). Those strains and the two other *Drosophila* species tested, *D. busckii* (derived from flies collected in South-East France in 2015) and *D. suzukii* (gift from Dr. R. Allemand, LBBE, University of Lyon 1), were reared under controlled laboratory conditions (150-200 eggs/40 ml fly medium; 25°C for *D. melanogaster* and 20°C for the two other fly species; 60% relative humidity; 12:12 light/dark cycle) on a high-protein/sugar-free fly medium (10% cornmeal, 10% yeast, 0% sugar). All the experiments were performed under these laboratory conditions.

### (b) Bacillus thuringiensis bioinsecticide product

Spores and Cry toxins of *Bt.* var. *kurstaki* strain SA-11 were from a commercial bioinsecticide product (Delfin^®^ wettable granules, Valent BioSciences, AMM 9200482, 32,000 UI/mg).

Viable spores were estimated at 5×10^7^ CFU/mg product by counting Colony Forming Units (CFUs) on LB agar, and this value remained stable during the timeframe of this study. For the experiments, suspensions of *Btk* bioinsecticide were prepared in Ringer buffer (NaCl 7.5g/l, NaHCO_3_ 0.1g/l, KCl 0.2g/l, CaCl2 0.2g/l, in distilled water) to reach the desired CFUs in 100 µl.

### (c) Oviposition choice test

Two-to-five day-old mated females (20 *D. melanogaster*, 30 *D. suzukii*, 30 *D. busckii*) were transferred to aerated plastic cages (Ø 10.5 cm, h 7.5 cm) containing two dishes (Ø 3 cm, ∼7 cm^2^, 1g of fly medium) diametrically opposed at the cage bottom. The test lasted 18 h for *D. melanogaster* and 24 h for *D. suzukii* and *D. busckii* which lay fewer eggs. To avoid confounding effects, cage orientation and location in the experimental chamber were randomized.

### (d) Oviposition in presence of Btk bioinsecticide

Flies were given the choice between a dish filled with fly medium mixed with a suspension of *Btk* bioinsecticide in Ringer buffer at a given concentration, and a control dish filled with fly medium mixed with the same volume of Ringer buffer (dose “0”). In control replicate cages, females were offered the choice between two dishes filled with fly medium mixed with Ringer buffer. Oviposition preference for *Btk* was calculated by dividing the number of eggs laid on the *Btk* substrate divided by the sum of eggs on the two substrates of the cage.

Oviposition preference equal to 0.5 indicates no preference or avoidance of the bioinsecticide; preference values above 0.5 indicate bioinsecticide appetitiveness, while values below 0.5 indicate bioinsecticide avoidance. Oviposition preference in control cages was the egg proportion on one of the two Ringer substrates. For each cage, the fly motivation for egg laying was assessed by summing the eggs laid on the two substrates.

Three *Btk* bioinsecticide doses previously described in ^[27]^ were used: 10^6^ CFU/g fly medium that has no effect on the *Drosophila* development and falls in the recommendation range (equivalent to the field application of 1.4×10^5^ CFU/cm^2^) and 10^8^ CFU/g and 10^9^ CFU/g which strongly alters *Drosophila* larval development (equivalent to the application of 1.4×10^7^ CFU/cm^2^ and 1.4×10^8^ CFU/cm^2^, respectively). The dynamics of egg laying over the 18-h choice test were explored with the *D. melanogaster* Canton-S strain by measuring the oviposition preference at 2 h, 4 h, and 18 h (endpoint) of choice test. Oviposition preference of *D. suzukii* and *D. busckii* was measured with the choice over 24 h between a Ringer control substrate and a substrate containing 10^9^ CFU/g of *Btk* bioinsecticide.

To disentangle the effects on the oviposition preference of *Btk* spores, toxin crystals and soluble toxins, from those of the commercial product additives, a 2×10^10^ CFU suspension of the bioinsecticide product was dialyzed to remove low molecular weight compounds.^[27]^ A fraction of the dialyzed suspension was centrifuged at 15,000 g, 15 min, 18°C to collect the pellet containing mainly spores and toxin crystals, and the supernatant containing toxin fragments and non-dialyzable compounds.^[27]^ The oviposition preference and motivation for egg laying of *Drosophila melanogaster* Canton-S females was assessed during 18 h when flies were offered the choice between a control Ringer substrate and a substrate containing the non-dialyzed bioinsecticide, the dialyzed bioinsecticide, the centrifugation pellet (all adjusted to 10^9^ CFU/g), the supernatant, or the PBS buffer used for dialysis.

### (e) Statistical analysis

Binomial data on oviposition preference were analysed with mixed-effects generalized linear models that included, when appropriate, the *D. melanogaster* strain, the *Btk* treatment (Ringer control, *Btk* bioinsecticide doses, dialysis and centrifugation fractions), the choice test duration and their two-way interactions as fixed factors. The replicate cage was included as random factor. Count data on egg-laying motivation were transformed into decimal logarithm values and analysed with mixed-effect models including the same fixed and random effects as described above (similar statistical results and biological conclusions were obtained with untransformed data). Significance of fixed effects and interactions was tested by model comparisons. Pairwise *post hoc* comparisons of each *Btk* dose with the no-*Btk* control and of each fly strain with Canton-S were performed. The deviation of the oviposition preference from a 50%-50% distribution of eggs on the two substrates was tested with t tests under the H0 hypothesis of a mean egg proportion of 0.5. The replicate number being relatively small, Wilcoxon tests with the same H0 hypothesis were performed and yielded similar biological conclusions. Statistical analyses were performed in R^[55]^ using the packages lme4^[56]^ and multcomp.^[57]^

## 3. Results

### (a) Drosophila melanogaster expressed a rapid, dose-dependent oviposition avoidance of Btk bioinsecticide

The presence of *Btk* bioinsecticide impacted the oviposition preference of *D. melanogaster* females over 18h compared to the controls without bioinsecticide, yet with varying amplitudes between fly strains (Figure 1; Table S1.1). Canton-S females laid eggs evenly when offered two control substrates, while they laid fewer eggs on *Btk* substrate when offered a choice between substrates with and without *Btk* (Table S1.1; significance of *post hoc* control-*Btk* dose pairwise comparisons in Figure 1). Their *Btk* avoidance increased with the bioinsecticide dose, and deviated significantly from the “neutral” preference of 0.5 at the two highest doses, 10^8^ and 10^9^ CFU/g (Table S1.1), dropping to 0.19 on average at 10^9^ CFU/g (95% confidence interval: 0.07 – 0.30). The oviposition preference of Nasrallah females also decreased with the increasing *Btk* dose (Figure 1; Table S1.1), dropping significantly below 0.5 only at 10^9^ CFU/g with a smaller amplitude than that of Canton-S females (0.27 on average, 95% CI: 0.12 – 0.41; Table S1.1). Similarly, the average preference of Sefra females was 0.29 at this dose (95% CI: 0.21 – 0.37), while the dose 10^6^ CFU/g was slightly appetitive (Figure 1; Table S1.1). The oviposition preference of the YW double mutant also decreased significantly below 0.5 at 10^9^ CFU/g but with a smaller amplitude (average preference of 0.37, 95% CI: 0.24 – 0.50) (Figure 1; Table S1.1). For all the four *D. melanogaster* strains, female motivation for egg laying in the presence of *Btk* bioinsecticide was similar to that without bioinsecticide and was similar between *Btk* doses (Figure 2; Table S1.1).

**Figure 1.**
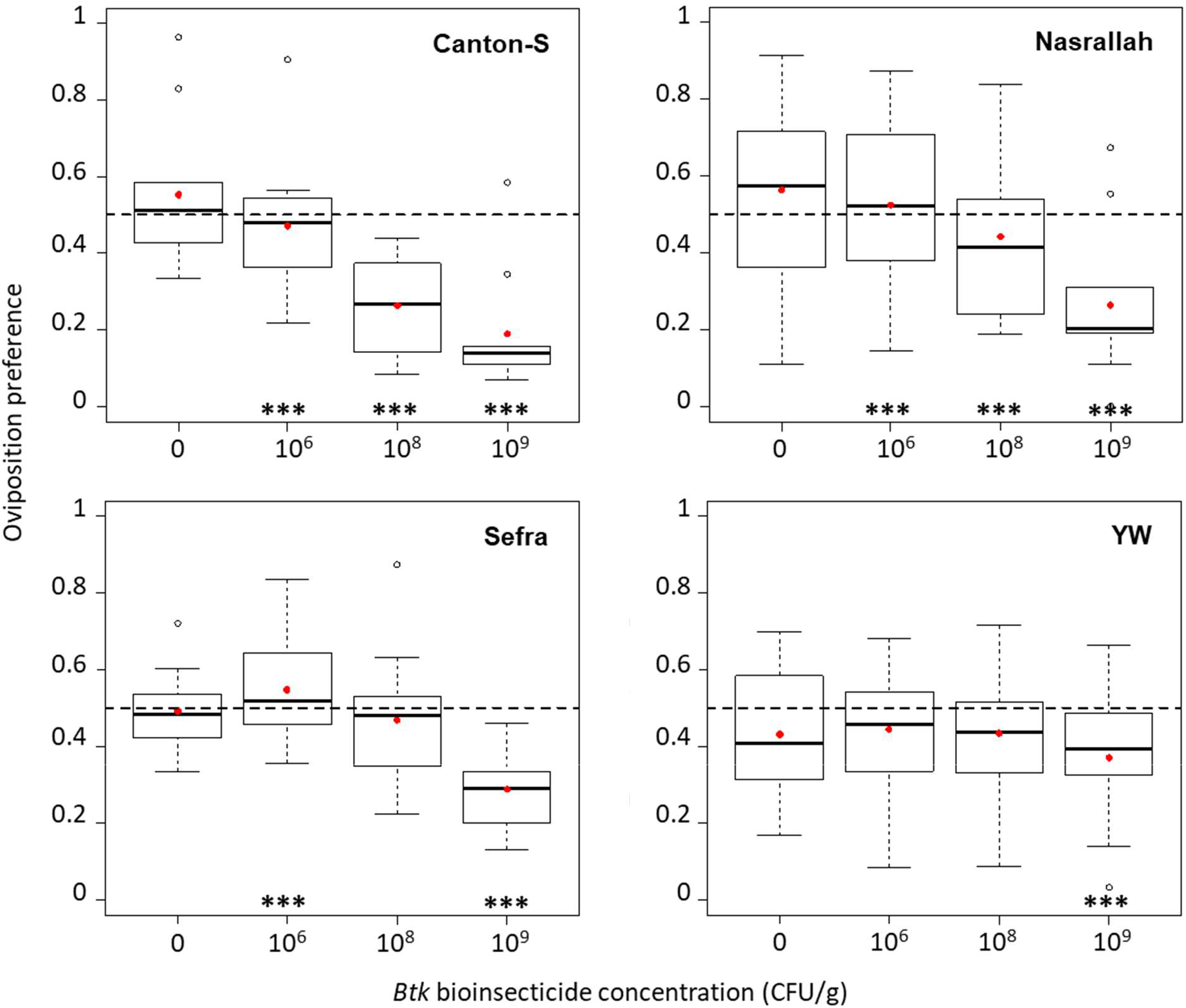
Female oviposition preference in the 18-hour choice test as the proportion of eggs laid on one food substrate (quartiles, median and mean preference in red points) of *Drosophila melanogaster* wild-type strains Canton-S, Nasrallah and Sefra, and the YW double mutant strain, with three doses of *Btk* bioinsecticide (10^6^, 10^8^, and 10^9^ CFU/g of fly medium) and the no-*Btk* Ringer control (0). Significance of *post hoc* pairwise comparisons of the control with each *Btk* dose: *** *P* < 0.0001. *N* = 10 replicate cages per treatment for each fly strain.

**Figure 2.**
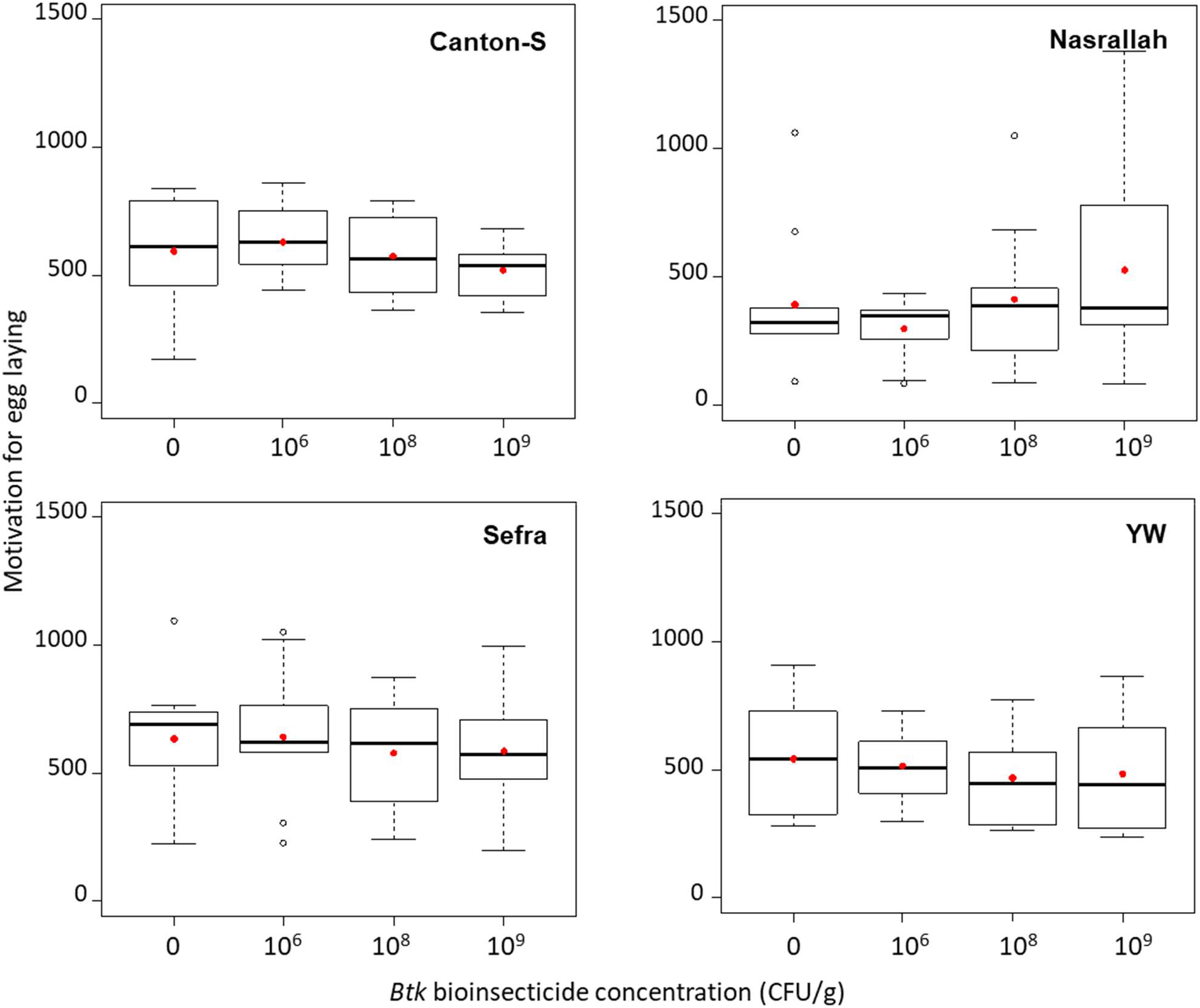
Fly motivation for egg laying during the 18-hout oviposition choice test as the total number of eggs laid on both food substrates offered (quartiles, median and mean of the total number of eggs in red points) of *Drosophila melanogaster* wild-type strains Canton-S, Nasrallah and Sefra, and the YW double mutant strain, with 3 doses of *Btk* bioinsecticide (10^6^, 10^8^, and 10^9^ CFU/g of fly medium) and the no-*Btk* Ringer control (0). *N* = 10 replicates cages per treatment for each fly strain.

Over the course of the 18-h choice test, the oviposition preference of the control Canton-S females unexposed to *Btk* bioinsecticide did not differ from the “neutral” preference 0.5 despite random variation between time points. In contrast, when offered the choice between a *Btk* substrate at 10^9^ CFU/g and a control substrate, the female preference for *Btk* was already below 0.5 at 2 h and further decreased at 4 h to remain down to ∼0.2 until the end of the choice test (Figure 3A; Table S1.2). The motivation of Canton-S females for egg laying evolved similarly and regardless of the choice they were offered (Figure 3B; Table S1.2).

**Figure 3.**
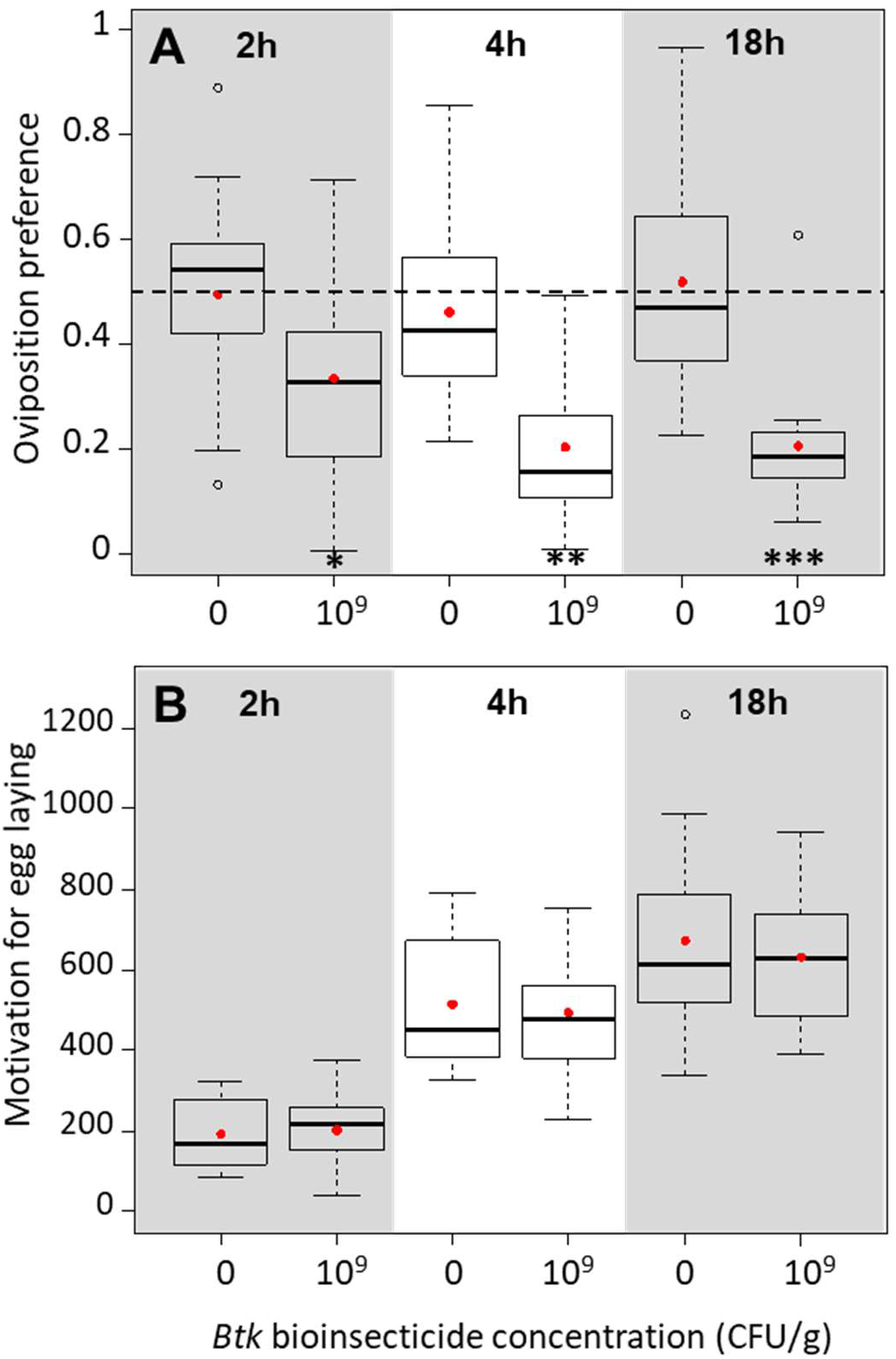
Dynamics of *Drosophila melanogaster* Canton-S female (A) oviposition preference as the proportion of eggs laid on one food substrate, and (B) motivation for egg laying as the total number of eggs laid on both food substrates (quartiles, median and mean per treatment in red points) recorded at 2h, 4h, and 18h in the oviposition choice test with 10^9^ CFU/g of *Btk* bioinsecticide and the no-*Btk* Ringer control (0). Significance of *post hoc* pairwise comparisons of the control with the *Btk* bioinsecticide: * *P* < 0.05, ** *P* < 0.01 *** *P* < 0.0001. *N* = 15 replicate cages per treatment and test duration.

### (b) All the Btk bioinsecticide fractions elicited the fly oviposition avoidance

While the preference after 18h of Canton-S females for both Ringer and PBS controls did not differ from 0.5 (Figure 4A, Table S2), females significantly avoided the dialyzed *Btk* suspension, the suspended pellet and the supernatant with a similar amplitude as the non-dialyzed *Btk* bioinsecticide at 10^9^ CFU/g (average preference of 0.30, 95% CI: 0.21 – 0.39; Figure 4A, Table S2). The female motivation for egg laying was similar across choice modalities (Figure 4B, Table S2).

**Figure 4.**
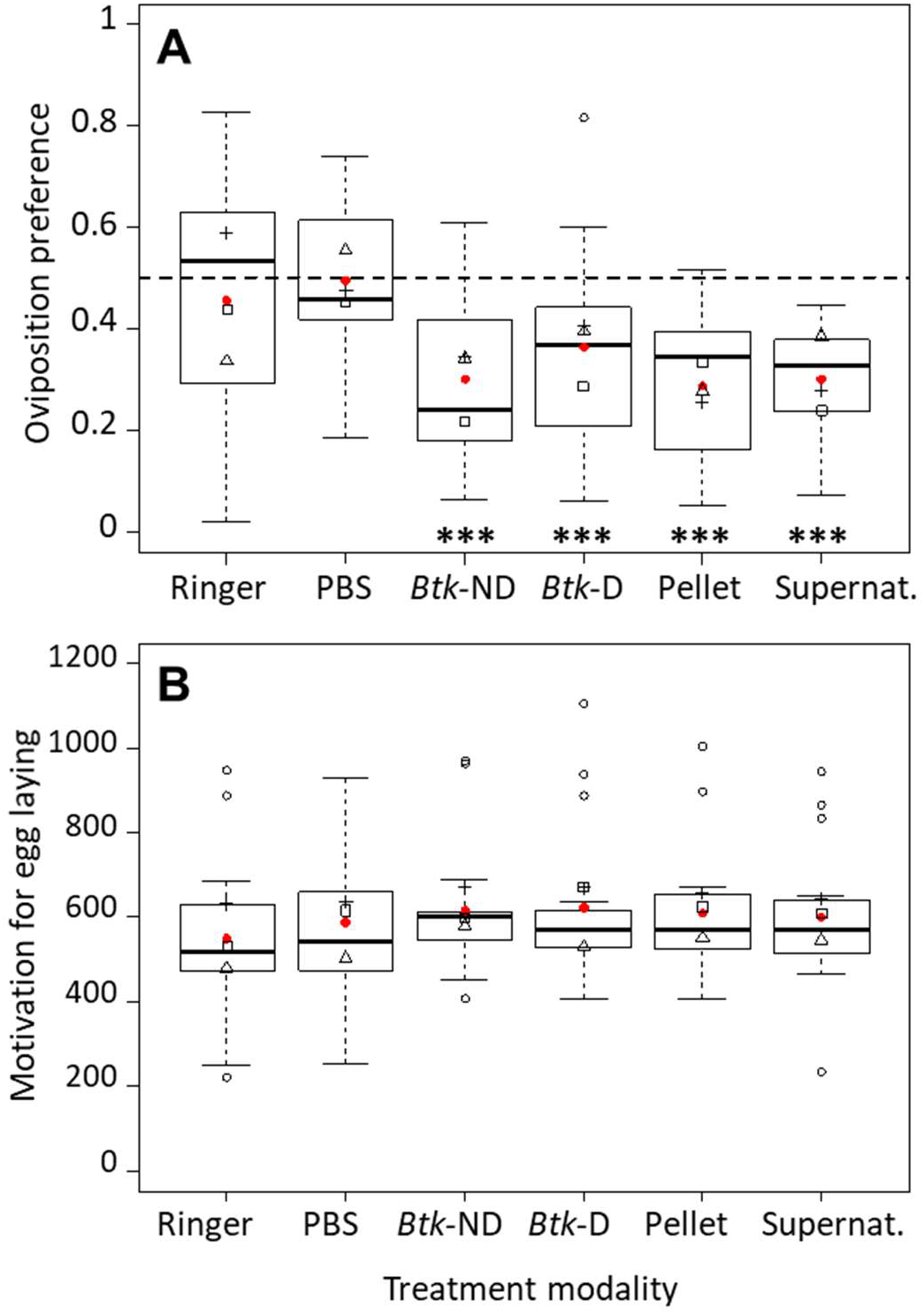
*Drosophila melanogaster* Canton-S female (A) oviposition preference as the proportion of eggs laid on one food substrate, and (B) motivation for egg laying as the total number of eggs laid on both food substrates (quartiles, median and mean per treatment in red points) in the 18-hour oviposition choice test with *Btk* bioinsecticide at 10^9^ CFU/g of fly medium (*Btk-*ND), dialyzed *Btk* bioinsecticide (*Btk*-D) and the pellet (Pellet) adjusted to the same concentration, the supernatant (Supernat.) after centrifugation, and the Ringer and PBS controls. Significance of *post hoc* pairwise comparisons of the Ringer control with each of the other treatment modalities: *** *P* < 0.001. *N* = 15 replicates cages per treatment.

### (c) The amplitude of fly avoidance of Btk bioinsecticide varied between species

Females of the invasive species *D. suzukii* strongly avoided *Btk* in the choice test: their oviposition preference dropped to 0.16 on average at 10^9^ CFU/g of *Btk* (95% CI: 0.11 – 0.21; Figure 5A, Table S3), the results being similar when including only cages with >15 eggs (Figure S4). *Drosophila busckii* females’ preference also dropped significantly to 0.38 on average in presence of 10^9^ CFU/g *Btk* (95% CI: 0.28 – 0.49; Figure 5C, Table S5). The female motivation for egg laying of the two species was independent of the choice they were offered (Figures 5B, D; Figure S4; Tables S3, S5).

**Figure 5.**
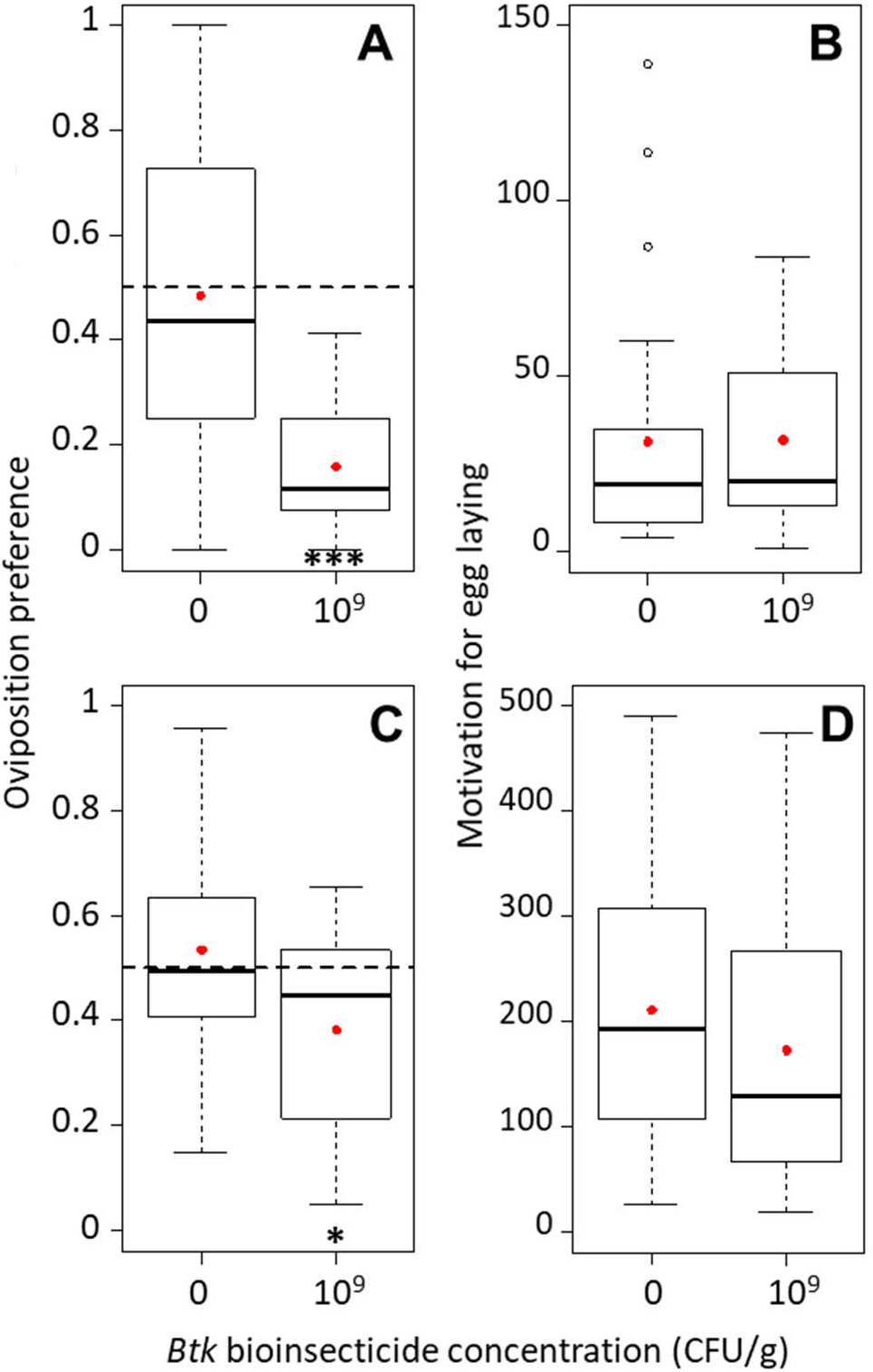
Drosophila suzukii *(A) and* Drosophila busckii (C) female oviposition preference as the proportion of eggs laid on one food substrate, and their respective motivation for egg laying (B, D) as the total number of eggs laid on both food substrates during the 24-hour oviposition choice test with *Btk* bioinsecticide at 10^9^ CFU/g and the no-*Btk* Ringer control (0) (quartiles, median and mean per treatment in red points). Significance of *post hoc* pairwise comparisons of the control with the *Btk* bioinsecticide: * *P* < 0.05 and *** *P* < 0.0001. *N* = 25 replicate cages per treatment for *D. suzukii* (all cages) and *N* = 15 replicate cages for *D. busckii*.

## 4. Discussion

When offered the choice between laying eggs on uncontaminated substrates or on *Btk* contaminated substrates, females of four strains of *Drosophila melanogaster* and of *D. busckii* and *D. suzukii* expressed avoidance of the *Btk* bioinsecticide. These oviposition responses were independent of confounding differences in the female motivation for egg laying. Since only non-ageing mated females were used during a short experimental period, this excludes the potential confounding effects of the female mating status and disturbance by male courtship, of sensory ageing impairing the ability to discriminate between food substrates, and of general ageing influencing the number of eggs laid.

*Drosophila melanogaster* strains avoided the bioinsecticide in a dose-dependent manner, and the three wild-type strains (Canton-S, Nasrallah, and Sefra) showed a stronger avoidance of the highest bioinsecticide dose than the YW double mutant strain. The smaller avoidance amplitude by YW females coincides with the fact that the yellow and white mutations affect the flies’ non-social and social behaviour and their ability to learn with olfactory cues.^[58–61]^ The invasive Asian species, *Drosophila suzukii*, exhibited a strong avoidance as the wild-type *D. melanogaster* Canton-S, although this species underwent an evolutionary shift in the bitter taste perception and oviposition preferences in the presence of microorganisms compared to other frugivorous *Drosophila* species.^[62,63]^ The third tested species, the opportunistic frugivorous *Drosophila busckii,* from the subgenus *Dorsilopha*, belongs to the *Drosophila* cosmopolitan guild of domestic species along with *D. melanogaster* and is specialized on vegetables.^[46]^ This species was the least avoidant, indicating that, although the bioinsecticide avoidance was general to our study, there was inter-species variability within the cosmopolitan guild.

*Drosophila melanogaster* behavioural avoidance of *Btk* bioinsecticide was detectable as early as 2 hours after the choice test onset, with increasing amplitude in the following few hours. The time scale of our results is consistent with previous reports of rapid learned avoidance towards pathogenic bacteria previously observed in *D. melanogaster*^[12]^ and *Caenorhabditis elegans.*^[13]^ The avoidance of the bioinsecticide may have started even earlier during the choice test, yet counting laid eggs does not allow a fine time resolution since a robust result requires substantial numbers of laid eggs. A video tracking of the fly positions in the cage might indicate whether females innately avoided *Btk* bioinsecticide (*i.e.* from the test onset) or were initially attracted to it and shifted their preference during the test, although *Drosophila* positional and oviposition preferences do not necessarily match.^[64]^ Nevertheless, our study showed that the expression of *Btk* bioinsecticide avoidance is rapid on a fly’s lifetime scale. The female decision-making for oviposition is a highly complex and dynamic trait that combines several parameters: the female’s genotype and experience of the oviposition substrates,^[12,65–67]^ the presence at oviposition sites of the male-derived aggregation pheromone transmitted to females during mating and emitted by recently mated females and of the deterring host marking pheromone,^[68–70]^ the social transmission of oviposition substrate preferences between females^[71–73]^ and of other information linked to substrate quality (presence of larvae and faeces),^[70,74,75]^ the presence of specific commensal microorganisms,^[63,76]^ the amplification of pheromone aggregation signal in infected flies by pathogenic bacteria^[77]^ and the group size.^[78]^ In addition, the texture of the oviposition substrate also plays an important role in the female oviposition decisions.^[63,79]^ In our study system, the bioinsecticide or Ringer buffer addition to the fly medium changed similarly the texture of the food substrate and did not change its pH.^[27]^

Behavioural avoidance was observed consistently for all the strains and species at the highest tested dose, 10^9^ CFU/g of *Btk* bioinsecticide, and at 10^8^ CFU/g for *D. melanogaster* Canton-S and Nasrallah. This mirrors the recent report of development alterations upon chronic exposure to these doses of *Btk* bioinsecticides, and the smaller bioinsecticide impacts on the emergence rates of *D. melanogaster* YW and *D. busckii* compared to other *D. melanogaster* strains and *D. suzukii*.^[27]^ While the dose 10^9^ CFU/g, which is 1,000 times above the manufacturer’s recommendations, seems unrealistic in the field, the dose 10^8^ CFU/g (equivalent to a field application of 1.4×10^7^ CFU/cm^2^)^[27]^ is reachable under current agriculture and horticulture practices where repeated applications are usually recommended (up to 8 authorized applications^[80,81]^ www.certiseurope.fr; www.certisusa.com). Indeed, *Bt* spores and toxins naturally persist and could accumulate in the field^[16,23,24,82]^ and bioinsecticide products contain protective compounds to lengthen their activity after field application.^[80,83]^ Very recently, doses close to 10^8^ CFU/g have been measured in honey bee matrices and flowers after the field application of the maximum recommended *Bt* bioinsecticide dose and concentrations up to 10^7^ CFU/g still persisted two days after application.^[84]^

Expression of behavioural avoidance toward *Btk* bioinsecticide was observed with the regular suspension, as well as with the dialyzed suspension and each of its fractions independently. This excludes a role in the avoidance of small molecular weight compounds of the formulation,^[85]^, and suggests that of spores, toxins, or residual bacterial fragments. Since *Bt* spores persist longer in the field than toxins,^[16,23,24]^ our results might indicate that the presence of spores in the environment may be sufficient to mediate bioinsecticide avoidance expression by non-target *Drosophila* females. Moreover, it was demonstrated that the nematode *C. elegans*^[86]^ and *D. melanogaster* males and females^[14]^ exhibit bacteria avoidance based on the presence of bacterial cell wall components. Here, the fractions of the bioinsecticide suspension after dialysis may contain cell wall components of the vegetative bacteria from the bioinsecticide manufacturing, which presence and role in the oviposition avoidance remain to be evaluated. When foraging, larvae and adult *Drosophila* naturally avoid specific harmful compounds or nutritionally unsuitable food based on the sensory perception of olfactory cues,^[87–89]^gustatory cues,^[62,90–92]^ or the physiological consequences of ingesting virulent bacteria.^[12]^ In our case, it seems unlikely that female avoidance of *Btk* bioinsecticides for oviposition relies only on olfactory cues, as this would likely result in stronger oviposition avoidance early during the test. The involvement of gustatory cues when assessing the oviposition site (e.g., bitter taste) and/or on physiological consequences of ingesting *Btk* bioinsecticide remains to be assessed.

From the point of view of females’ offspring, the oviposition avoidance of *Btk* bioinsecticide alleviates the cost of developing under chronic bioinsecticide exposure. Indeed, the growth and gut physiology of *D. melanogaster* larvae is already dramatically disturbed at 5×10^7^ CFU/g of *Btk* bioinsecticide.^[28]^ In addition, emergence rates of *D. melanogaster* strains developing on 10^8^ CFU/g of *Btk* bioinsecticide dropped by up to 81% compared to unexposed controls and groups exposed to only 10^6^ CFU/g.^[27]^ The development success was even null at the highest tested dose of *Btk* bioinsecticide, 10^9^ CFU/g.^[27]^ Avoidance of *Btk* bioinsecticide by females while searching for oviposition sites would thus increase their inclusive fitness, since more of their progeny would have a chance to develop and reach the adult stage and reproduce. Given that *Drosophila* females both feed and lay eggs on food substrates, the avoidance of *Btk* contaminated oviposition sites would also reduce the adult fly exposure to bioinsecticide, although adults do not seem to be impacted.^[27]^

From an ecological point of view, variations in avoidance amplitude between *D. melanogaster* genotypes and *Drosophila* species may modify their competitive interactions in *Btk*-treated areas. Interestingly, variations in avoidance strength have already been observed for carbon dioxide and other odorants indicating the stage of the fruit ripeness. These observations reflect the biological differences in feeding, mating, and oviposition between *Drosophila* species specialized on overripe fruits (several genotypes of *D. melanogaster*, *D. yakuba*, *D. pseudobscura*, *D. virilis*) and *D. suzukii* specialized on ripening fruits.^[87,88]^ In our study, the smaller amplitude of *Btk* bioinsecticide avoidance of the opportunistic *D. busckii,* combined with its lower developmental susceptibility to chronic bioinsecticide exposure^[27]^ suggest that *Btk* applications might not dramatically affect the field presence of this species in the *Drosophila* community. By contrast, the high susceptibility of *D. suzukii* to developmental alterations upon chronic exposure to bioinsecticide^[26,27]^ combined with the females’ amplitude of oviposition avoidance, suggest that developmental alterations could be alleviated by avoidance of *Btk-*treated areas. Despite the fact that *D. melanogaster* and *D. suzukii* have different niche specializations, their potential indirect interactions would be displaced mostly to untreated areas since both species avoid strongly the bioinsecticide for egg laying. The population dynamics of their natural enemies (predators and parasites) would be impacted by the changes in the location of their prey/host populations, in addition to be impacted directly by the bioinsecticide as previously reported for two species of *D. melanogaster* parasitoids.^[93]^ Indirectly, our results further suggest that *Btk* bioinsecticide application may be useful as a repellent to *D. suzukii* in orchards and gardening, but it may not be an efficient tool to control populations of this invasive fly and comes with side-effects on other non-target species.

In summary, females of several *Drosophila* species and genotypes expressed oviposition avoidance of food substrates contaminated with *Btk* bioinsecticide. The avoidance appeared rapidly after the onset of choice tests, for all the fractions of the bioinsecticide suspension and was independent of female motivation for egg laying. Our study extends the assessment of *Btk* bioinsecticide chronic effects previously reported in multiple *Drosophila* species to behavioural aspects, and highlights the need for multi-component assessments (development, physiology, life history, behaviour) of the potential effects of bioinsecticides on non-target invertebrates.

## Acknowledgements

We would like to thank Ms Jingru Li, Dorine Maertens and Yao Zhou for help with preliminary experiments, as well as Laurent Kremmer and Christian Rebuf for help with fly stocks.

## Author contributions

AB: conceptualization, data curation, formal analysis, investigation, methodology, writing-original draft, writing-review and editing; JLG: conceptualization, writing-review and editing; MP: funding acquisition, writing-review and editing.

