## Supplementary material for "*Bacillus thuringiensis* bioinsecticide influences *Drosophila* oviposition decision"

**Contents**

S1. Detailed statistical analysis of *D. melanogaster* preference and motivation for oviposition

S2. Detailed statistical analysis of *D. melanogaster* preference and motivation for oviposition in presence of the *Btk* bioinsecticide components

S3. Detailed statistical analysis of *D. suzukii* preference and motivation for oviposition

S4. *Drosophila suzukii* preference and motivation for oviposition including only part of the data

S5. Detailed statistical analysis of *D. busckii* preference and motivation for oviposition

### S1. Detailed statistical analysis of *D. melanogaster* preference and motivation for oviposition

The tables below summarize the results of the statistical analyses carried out to test the effect of the *Btk* bioinsecticide dose, *D. melanogaster* strain, their two-way interactions, the time point and the replicate cage as random effect. Results of *post hoc* tests are shown for significant fixed effects.

**Table S1.1.** Statistical analysis of the oviposition preference (proportion of eggs laid on the *Btk* substrate or on one of the control substrates) and motivation for egg laying (total number of eggs laid on both substrates) of the four *D. melanogaster* strains (CantonS, Nasrallah, Sefra and YW) after 18h oviposition choice test.

| **Source of variation** | ***df*** | $\chi^2$ | ***P*** |
|---|---|---|---|
| ***(a) Oviposition preference*** | | | |
| Fly strain × *Btk* dose | 9 | 1355 | **<0.0001** |
| * CantonS – Nasrallah: -0.028 ± 0.042, $z$ = -0.65, $P$ = 0.86 | | | |
| * CantonS – Sefra: -0.36 ± 0.04, $z$ = -9.82, $P$ < 0.0001 | | | |
| * CantonS – YW: -0.50 ± 0.04, $z$ = -13.04, $P$ < 0.0001 | | | |
| *Btk* dose effect per fly strain: | | | |
| * CantonS | 3 | 2445 | **<0.0001** |
| Pairwise comparisons with control: no *Btk* – $10^6$: -0.45 ± 0.04, $z$ = -12.11, $P$ < 0.0001; no *Btk* - $10^8$: -1.38 ± 0.04, $z$ = -33.84, $P$ < 0.0001; no *Btk* – $10^9$: -1.84 ± 0.05, $z$ = -40.35, $P$ < 0.0001 | | | |
| Difference with neutral preference 0.5: no *Btk*: $t$ = 0.85, $df$ = 9, $P$ = 0.42; $10^6$: $t$ = -0.45, $df$ = 9, $P$ = 0.66; $10^8$: $t$ = -5.99, $df$ = 9, $P$ = 0.0002; $10^9$: $t$ = -6.22, $df$ = 9, $P$ = 0.00015 | | | |
| * Nasrallah | 3 | 1029 | **<0.0001** |
| Pairwise comparisons with control: no *Btk* – $10^6$: -0.22 ± 0.05, $z$ = -4.20, $P$ < 0.0001; no *Btk* - $10^8$: -0.42 ± 0.05, $z$ = -8.93, $P$ < 0.0001; no *Btk* – $10^9$: -1.40 ± 0.05, $z$ = -28.89, $P$ < 0.0001 | | | |
| Difference with neutral preference 0.5: no *Btk*: $t$ = 0.80, $df$ = 9, $P$ = 0.44; $10^6$: $t$ = 0.33, $df$ = 9, $P$ = 0.75; $10^8$: $t$ = -0.81, $df$ = 9, $P$ = 0.44; $10^9$: $t$ = -3.66, $df$ = 9, $P$ = 0.0052 | | | |
| * Sefra | 3 | 1011 | **<0.0001** |
| Pairwise comparisons with control: no *Btk* – $10^6$: 0.41 ± 0.04, $z$ = 11.41, $P$ < 0.0001; no *Btk* - $10^8$: -0.01 ± 0.04, $z$ = -0.29, $P$ = 0.99; no *Btk* – $10^9$: -0.76 ± 0.04, $z$ = -20.03, $P$ < 0.0001 | | | |
| Difference with neutral preference 0.5: no *Btk*: $t$ = -0.20, $df$ = 9, $P$ = 0.85; $10^6$: $t$ = 1.03, $df$ = 9, $P$ = 0.33; $10^8$: $t$ = -0.48, $df$ = 9, $P$ = 0.64; $10^9$: $t$ = -6.27, $df$ = 9, $P$ = 0.00015 | | | |
| * YW | 3 | 75.9 | **<0.0001** |
| Pairwise comparisons with control: no *Btk* – $10^6$: 0.08 ± 0.04, $z$ = 1.99, $P$ = 0.12; no *Btk* - $10^8$: 0.05 ± 0.04, $z$ = 1.27, $P$ = 0.45; no *Btk* – $10^9$: -0.25 ± 0.04, $z$ = -6.0, $P$ < 0.001 | | | |
| Difference with neutral preference 0.5: no *Btk*: $t$ = -1.15, $df$ = 9, $P$ = 0.28; $10^6$: $t$ = -1.01, $df$ = 9, $P$ = 0.34; $10^8$: $t$ = -1.07, $df$ = 9, $P$ = 0.31; $10^9$: $t$ = -2.25, $df$ = 9, $P$ = 0.051 | | | |
| ***(b) Motivation for egg laying (*Btk *dose effect for each fly strain)*** | | | |
| * CantonS | 3 | 3.87 | 0.28 |
| * Nasrallah | 3 | 1.43 | 0.70 |
| * Sefra | 3 | 5.40 | 0.15 |
| * YW | 3 | 1.01 | 0.80 |

**Table S1.2.** Statistical analysis of the dynamics of the oviposition preference (proportion of eggs laid on the *Btk* substrate or on one of the control substrates) and motivation for egg laying (total number of eggs laid on both substrates) of *D. melanogaster* CantonS females during the 18h oviposition choice test.

| **Source of variation** | **df** | $\chi^2$ | **P** |
|---|---|---|---|
| ***(a) Oviposition preference*** | | | |
| *Btk* dose × Time | 2 | 112.1 | **<0.0001** |
| Pairwise comparisons of *Btk*-control treatments at each time point: 2h: $t = 2.16$, $df = 28$, $P = 0.04$; 4h: $t = 4.19$, $df = 27$, $P = 0.0003$; 18h: $t = 4.95$, $df = 23$, $P < 0.0001$ | | | |
| Time effect per *Btk* dose: | | | |
| * no-*Btk* control | 2 | 48.0 | **<0.0001** |
| Pairwise comparisons: 2h – 4h: -0.08 ± 0.05, $z = -1.72$, $P = 0.19$; 2h – 18h: 0.14 ± 0.04, $z = 3.1$, $P = 0.005$; 4h – 18h: 0.22 ± 0.03, $z = 6.86$, $P < 0.0001$ | | | |
| Difference with neutral preference 0.5: 2h: $t = -0.06$, $df = 14$, $P = 0.96$; 4h: $t = -0.79$, $df = 14$, $P = 0.45$; 18h: $t = 0.36$, $df = 14$, $P = 0.73$ | | | |
| * *Btk* $10^9$ CFU/g | 2 | 138.2 | **<0.0001** |
| Pairwise comparisons: 2h – 4h: -0.52 ± 0.05, $z = -10.4$, $P < 0.0001$; 2h – 18h: -0.55 ± 0.05, $z = -11.4$, $P < 0.0001$; 4h – 18h: -0.04 ± 0.04, $z = -0.92$, $P = 0.63$ | | | |
| Difference with neutral preference 0.5: 2h: $t = -3.09$, $df = 14$, $P = 0.008$; 4h: $t = -7.52$, $df = 14$, $P < 0.0001$; 18h: $t = -9.03$, $df = 14$, $P < 0.0001$ | | | |
| ***(b) Motivation for egg laying*** | | | |
| *Btk* dose | 1 | 0.10 | 0.75 |
| Time | 2 | 95.2 | **<0.0001** |
| *Btk* dose × Time | 2 | 0.07 | 0.97 |

**S2. Detailed statistical analysis of *D. melanogaster* preference and motivation for oviposition in presence of the *Btk* bioinsecticide components**

The table below summarizes the results of the statistical analyses carried out to test the effect of different components of the *Btk* bioinsecticide as a fixed effect, and the replicate cage as a random effect. Results of *post hoc* tests are indicated for significant fixed effects.

**Table S2.** Statistical analysis of the oviposition preference (proportion of eggs laid on the *Btk*/fraction substrate or on one of the control substrates) and motivation for egg laying (total number of eggs laid on both substrates) of female *D. melanogaster* CantonS when offered the choice between a no-*Btk* control substrate and one of the modalities (Ringer, PBS, non-dialyzed *Btk* (*Btk*-ND), dialyzed *Btk* (*Btk*-D), Pellet, Supernatant).

| **Source of variation** | **df** | $\chi^2$ | **P** |
|---|---|---|---|
| ***(a) Oviposition preference*** | | | |
| Treatment: | 5 | 1844 | **<0.0001** |
| Pairwise comparisons: Ringer – PBS: -0.06 ± 0.03, *z* = -1.99, *P* = 0.17; Ringer – *Btk*-ND: -0.84 ± 0.03, *z* = -25.9, *P* < 0.001; Ringer – *Btk*-D: -0.60 ± 0.03, *z* = -18.9, *P* < 0.001; Ringer – Pellet: -0.99 ± 0.03, *z* = -29.9, *P* < 0.001; Ringer – Supernatant: -0.91 ± 0.03, *z* = -27.4, *P* < 0.001 | | | |
| Difference with neutral preference 0.5: Ringer: *t* = -0.67, *df* = 13, *P* = 0.52; PBS: *t* = -0.13, *df* = 14, *P* = 0.90; *Btk*-ND: *t* = -4.84, *df* = 14, *P* = 0.0003; *Btk*-D: *t* = -2.52, *df* = 14, *P* = 0.02; Pellet: *t* = -5.80, *df* = 14, *P* < 0.0001; Supernatant: *t* = -7.39, *df* = 14, *P* < 0.0001 | | | |
| ***(b) Motivation for egg laying*** | | | |
| Treatment | 5 | 4.72 | 0.45 |

**S3. Detailed statistical analysis of *D. suzukii* preference and motivation for oviposition**

The table below summarizes the results of the statistical analyses carried out to test the effect of the presence of *Btk* bioinsecticide as a fixed effect, and the replicate cage as a random effect. Results of *post hoc* tests are indicated for significant fixed effects.

**Table S2.** Statistical analysis of the oviposition preference (proportion of eggs laid on the *Btk* substrate or on one of the control substrates) and motivation for egg laying (total number of eggs laid on both substrates) of female *D. suzukii* when offered the choice between a no-*Btk* control substrate and a *Btk* substrate at $10^9$ CFU/g.

| **Source of variation** | **df** | $\chi^2$ | **P** |
|---|---|---|---|
| | ***(a) Oviposition preference*** | | |
| Treatment: | 1 | 162.8 | **<0.0001** |
| Difference with neutral preference 0.5: Ringer: $t$ = -0.25, $df$ = 24, $P$ = 0.81; $10^9$: $t$ = -14.5, $df$ = 24, $P$ < 0.0001 | | | |
| | ***(b) Motivation for egg laying*** | | |
| Treatment | 1 | 0.41 | 0.52 |

**S4. *Drosophila suzukii* preference and motivation for oviposition including only part of the data**

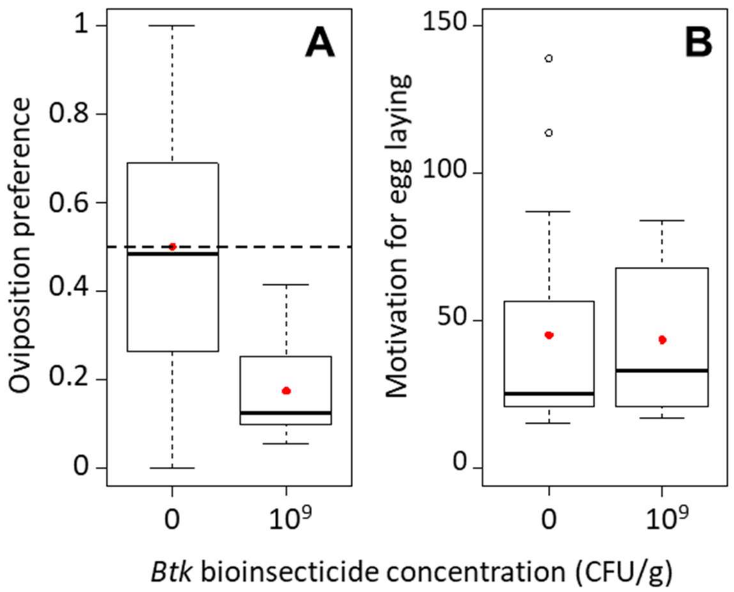


**Figure S4.** *D. suzukii* female (A) preference as the egg proportion on one food substrate (either the control substrate or the *Btk* substrate), and (B) motivation for egg laying as the total number of eggs on the two food substrates (quartiles and median, and mean per treatment as red lozenges) in the 24-hour oviposition choice test with *Btk* bioinsecticide at $10^9$ CFU/g and no-*Btk* Ringer control (0), including only the replicate cage with ≥ 15 eggs. *N* = 16 replicate cages for Ringer control and *N* = 16 replicate cages for *Btk* $10^9$ CFU/g.

**S5. Detailed statistical analysis of *D. busckii* preference and motivation for oviposition**

The table below summarizes the results of the statistical analyses carried out to test the effect of the presence of *Btk* bioinsecticide as a fixed effect, and the replicate cage as a random effect. Results of *post hoc* tests are indicated for significant fixed effects.

**Table S2.** Statistical analysis of the oviposition preference (proportion of eggs laid on the *Btk* substrate or on one control substrate) and motivation for egg laying (total number of eggs laid on both substrates) of female *D. busckii* when offered the choice between a no-*Btk* control substrate and a *Btk* substrate at $10^9$ CFU/g.

| **Source of variation** | **df** | $\chi^2$ | **P** |
|---|---|---|---|
| ***(a) Oviposition preference*** | | | |
| Treatment: | 1 | 137.2 | **<0.0001** |
| Difference with neutral preference 0.5: Ringer: $t = 0.57$, $df = 14$, $P = 0.58$; $10^9$: $t = -2.40$, $df = 14$, $P = 0.03$ | | | |
| ***(b) Motivation for egg laying*** | | | |
| Treatment | 1 | 1.27 | 0.26 |
